# Serial Dependence operates on categorical rather than physical representations: evidence from behavior and EEG

**DOI:** 10.64898/2026.01.22.700534

**Authors:** Pierre Costa, Thérèse Collins

**Affiliations:** Université Paris Cité, CNRS, Integrative Neuroscience and Cognition Center, F-75006 Paris, France

## Abstract

Our environment is constantly changing but our perception is stable. This stability may result from the integration of successive visual images that smooths over spurious variations in input. Such a process manifests in a behavioral phenomenon called Serial Dependence (SD), an attractive bias between successive stimuli. Here we questioned whether SD originates from physical stimulus representations or at a higher level, that of categorical perception. We constructed face morphs from three prototypes of the same face expressing anger, fear and sadness. We measured the perceived similarity between each pair of morphs with an odd-one-out experiment, to build a behavioral representational similarity matrix. Emotional expressions were perceived categorically. Subjects then reproduced the perceived facial expression of successively presented morphs by adjusting a response cue while their cerebral activity was recorded with EEG. We showed attractive serial dependence tuned to perceptual distance rather than physical similarity. We used representational similarity analysis to discover the geometry of the mental representation of faces, to capture both the information represented in a neural signal and its format. EEG representational similarity analyses showed that neural responses encoded emotional ambiguity rather than category membership, with near-prototypical faces eliciting similar patterns across emotions. Models combining feature tuning with categorical structure best accounted for behavioral biases. Neural signals also carried traces of past stimuli. Stimuli for which memory-based representations diverged most from sensory representations showed the strongest attractive serial dependence.

## INTRODUCTION

The outside world is in constant flux, from object movement to changes in how objects reflect light. Our reception of this information is also highly variable, since our own eye movements and blinks introduce additional sources of instability into retinal inputs. Despite this variability, visual perception is stable over time. This stability may result from the integration of consecutive visual images that smooths over spurious variations in input. If this is true, then the perception of the world that surrounds us is caused not only by current sensory input but also by the recent past. Such a process may manifest in a behavioral phenomenon called serial dependence (SD), an attractive bias between consecutive stimuli. This effect of the past, measurable by errors in judgements about the current stimulus, might be beneficial to the perceptual system given the natural statistics of the world, which is highly auto-correlated (Dong & Atick, 1995). An effect of the recent past on current perceptual experience has long been reported in the literature (Brascamp et al., 2008; Fecteau & Munoz, 2003), but the effect was termed serial dependence in 2014 (Cicchini et al., 2014; Fischer & Whitney, 2014) and has since then received renewed interest (Manassi et al., 2023; Pascucci et al., 2023). There is a debate in the literature about the level of processing at which SD occurs. Schematically, SD could be either low-level, related to a physical aspect of the stimulus or its features, or high-level, related to a memory or post-perceptual trace of the stimulus. Because stimuli and responses are often highly correlated, especially in tasks like orientation matching (often used to quantify SD), it is sometimes difficult to dissociate the influence of a previous response or decision from the influence of a previous stimulus. Nevertheless, some studies have shown that it is past decisions, rather than past stimuli, that exert an attractive effect on subsequent trials, i.e. observers tend to repeat the same response (Fritsche et al., 2017). The effect of past stimuli, according to some (but not all) studies, may be repulsive, as in the well-known adaptation effects (Fritsche et al., 2017). In fact, the effect of previous stimuli can be repulsive or attractive depending on stimulus duration and task (Can & Collins, 2025; Cicchini et al., 2017). Other studies have shown that SD is a perceptual effect, that current stimuli actually *look* like previous stimuli (Cicchini et al., 2017; Collins, 2020), suggesting that the site of SD is low-level or perceptual.

To investigate the level of visual processing at which SD occurs, we leveraged two well-known effects. The first is categorical perception. We used categorical perception as a tool to investigate whether SD occurs at the level of the mental representation of physical features, or of categorical membership. The perception of color is a classic example of categorical perception (Bornstein and Korda, 1984), as is the perception of facial emotional expression (Etcoff and Magee, 1992). We created face morphs that varied along a continuous feature space, but were perceived as categories: morphs tended to be perceived as more similar when they belonged to the same emotion category than when they did not. We evaluated the perceived similarity of each pair of morphs.

The second effect that we leveraged to investigate the level of visual processing at which SD occurs was its feature tuning. The influence of a stimulus on the subsequent stimulus is maximal for small inter-stimuli distances in feature space, and decreases when that distance is too large (Manassi et al., 2023). This feature-tuning property of SD allowed us to contrast whether the feature space relevant for SD was best defined as a physical continuum, or whether it was categorical. If SD operates on a categorically-defined representation of space, this suggests that the mechanisms behind SD operate at high, or at least intermediate, levels of visual processing.

We gathered behavioral and encephalographic (EEG) responses of observers viewing and judging the emotional expressions of faces.

Our analytical approach was representational similarity analysis (RSA), which abstracts away from specific signals to discover the geometry of a representation, and is a useful intermediate level of description because it captures both the information represented in a signal and its format (Kriegeskorte, 2008). RSA is thus the ideal tool for comparing behavior, neural activity, and theoretical models, which we do here. The perceptual similarity between faces with different emotional expressions allowed us to determine the geometry of the mental representation. The similarity between the neural activity evoked by different faces allowed us to determine the geometry of the neural representation. Because similarities are expressed as a matrix, the correlation between mental and neural representations can easily be ascertained, and we could thus compare their respective geometries, and compare them to our theoretical models. These models expressed low-level physical similarities between the faces, as well as several high-level characteristics.

We also applied RSA to examine the representations that underscore serial dependence. We examined how the response to one morph depended on the preceding morph. We examined the similarity between morph pairs by correlating the neural response evoked by one of the morphs in the pair with the neural response evoked by all morphs preceded by the other morph in the pair. This gave us a neural matrix expressing the similarity between a perceived morph (i.e. a sensory representation) and the past representation of another morph (i.e. a memory representation). We also adopted the classic analytical approach to serial dependence, which quantifies response errors as a function of the difference between two consecutive stimuli.

Serial dependence is a combination of past and current inputs to generate the current percept, and has often been framed within a Bayesian framework (Cicchini et al., 2023; Van Bergen & Jehee, 2019). A crucial question is which aspects of the past are integrated with which aspects of the present. In other words, what are the source and site of SD. Some studies have shown that priors that carry contextual information, and thus take their source in high-level, post-sensory representations of previous stimuli (Cicchini et al., 2021; Sheehan et al., 2024). In line with these findings, we hypothesized that SD would be modulated by the categorical perception of facial expression. Thus, we expected to find that SD was feature-tuned, and that this tuning was in categorical coordinates rather than physical. Moreover, we expected that the representational geometry that best matches both behavioral SD and the underlying neural representations would express high level aspects of the stimuli. If SD optimizes processing resources and behavior to the temporal structure of the natural environment, and if our hypotheses are verified, it would suggest that this optimization occurs at the level of meaningful, post-perceptual representations of events.

## METHODS

### Preliminary experiment

The goal of the preliminary experiment was to build a scale of facial expressions. We aimed to characterize the degree of perceived similarity between facial emotional expressions drawn from our set of stimuli.

### Participants

The preliminary experiment was conducted online with testable.org. We recruited 36 participants from their participant pool (15 female, 19-54 years old, M=34, SD=8.60).

### Stimuli

Stimuli were generated from three prototypes of the same face (a black-and-white Caucasian male face from the Radboud Face Database (RaFD); Langner et al., 2010; Figure 1a) expressing anger, fear and sadness, and 48 morphs between each prototype, yielding a total of 147 face stimuli. The morphs were generated using Sqirlz Morph 2.0 software. Points of interest on each prototype were hand-labeled. Morphing was then performed by a linear variation of the percentage of each of two prototypes (e.g. 25% anger and 75% fear would be one fourth of the way from fear to anger through the corresponding points of interest on each of those prototypes). Images were then cropped to an oval to remove the hairline, and a low-pass filter with a cutoff frequency of 17.5 cycles per degree in the Fourier domain was applied to each stimulus. Participants were asked to position themselves at arm’s length from their monitor and try to maintain this viewing distance throughout the experiment. On each trial, three faces were presented simultaneously, equidistantly from each other at 10 o’clock (320 pixels from center), 2 o’clock (320 pixels from center) and 6 o’clock (150 pixels from center). The size of each stimulus was 252 by 315 pixels.

**Figure 1.**
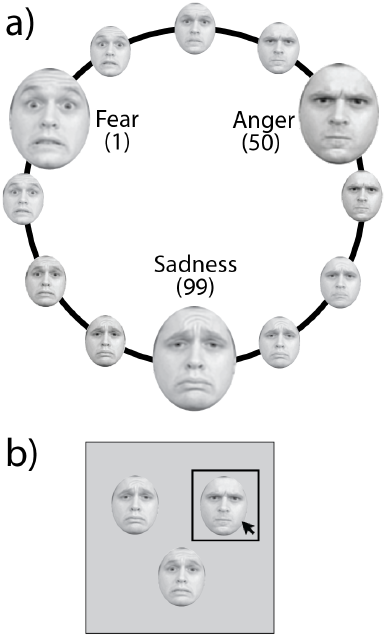
a) Stimuli were faces morphed between three prototypes (fear, anger, sadness). Three examples are shown but there were 48 morphs between each prototype in the experiments. Prototypes were taken from the Radboud Face Database (RaFD), morphs between prototypes were created for the purposes of the current experiment b) Preliminary experiment: triplet odd-one-out task. Participants clicked on the face that least resembled the two others.

### Procedure

Participants selected which of the three faces was the odd one out on each trial by clicking on it (Figure 1b). We did not provide explicit instructions as to the particular feature participants should use to decide. With 147 morphs, there are over three million possible different triplets. Collecting data over the entire set was impossible, so we pseudo-randomly sampled from the full list of possible triplets such as to gather similarity scores for short stimulus distances in finer detail than long distances. This enabled us to map our similarity matrix more efficiently with fewer trials. Participants went through a practice phase of three trials with feedback (which could be rerun as much as necessary). The test phase consisted of 500 trials. Each trial lasted until the participant responded. The whole experiment lasted approximately thirty minutes.

### Data analyses

Our hypothesis was that perception would be categorical: greater perceived similarity between faces belonging to the same category than between faces belonging to different categories, for a given physical difference. We used two different methods to test this hypothesis.

To characterize the perceived similarity between each pair of morphs, a similarity index was defined as the number of trials in which the pair was presented together but neither was selected as the odd one out, divided by the total number of trials that included the pair. For all the pairs that were not sampled, we used interpolation to infer the value of the perceived similarity (using the same method of interpolation and smoothing as in for the main experiment, see below). This allowed us to create a perceptual similarity matrix of size 147 by 147. To characterize the geometry of this matrix, we correlated it with theoretical models (see below). Our hypothesis here predicted that the perceptual similarity matrix would correlate the most with the categorical model.

Second, we labeled each morph as belonging to the category (fear, anger or sadness) of the predominant prototype (e.g. 10% anger and 90% fear was labeled as belonging to the fear category; there were no 50/50 stimuli). We grouped our data into two conditions: when a pair of stimuli belonged to the same category (within-category) versus when they did not (between-categories). The actual distance between morphs (in morph units) overlapped between the two conditions. We then examined whether perceived similarity depended on morph distance for each condition, hypothesizing that similarity would decrease with increasing morph distance, but that within-category similarity would always be higher than between-category similarity. We fitted a weighted exponential function to each condition (within-versus between-category) and performed permutation tests to check for a significant difference in intercepts. Across 1,000 iterations, we shuffled the mapping between perceived similarity and morph distance, refitted exponential functions, and derived a null distribution of intercept differences. Significance was set at alpha=0.05 (for this analysis and throughout).

### Main experiment

The goal of the main experiment was to measure serial dependence of facial expression. This experiment took place in the laboratory. The methods detailed below are inspired from Liberman et al. (2018).

### Participants

59 observers participated in the experiment. They reported no visual or neurological deficits and had normal or corrected-to-normal vision. 31 of them participated while their electroencephalographic (EEG) activity was recorded (19 female, 18-40 years old, M=26, SD=6.75). This number was determined by a stop rule. The 28 remaining participants provided only behavioral responses (19 female, 19-39 years old, M=27, SD=5.88). This number was determined by the amount of data necessary to fill in the behavioral serial dependence matrix. All participants provided written informed consent before taking part in the study. The study was approved by the French ethics committee (Comité de Protection des Personnes Ouest IV-Nantes) and followed the ethical guidelines of the Declaration of Helsinki (2013).

### Stimuli and Procedure

Participants completed an adjustment task in which they reproduced facial expressions by adjusting a response face to match a target expression (Figure 2).

**Figure 2.**
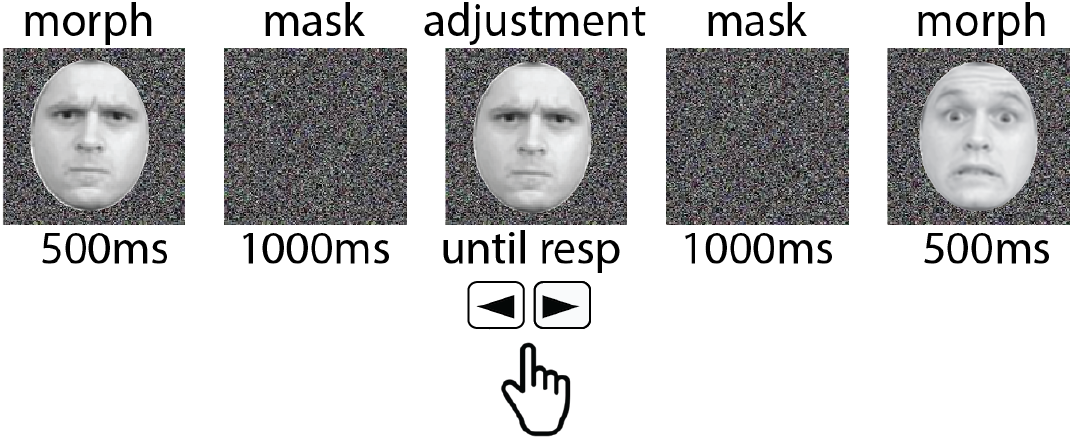
Task in the main experiment. A morph was presented for 500-ms, followed by a 1000-ms noise mask, followed by an adjustment face that participants could manipulate to match the remembered face. After giving their response, a 1000-ms noise mask was followed by a new morph. Prototypes were taken from the Radboud Face Database (RaFD).

We used the same morph stimuli as in the preliminary experiment. Each morph subtended 7 by 9 degrees of visual angle (dva) and was presented at screen center on a noise background. The experiment was programmed in MATLAB (The MathWorks, Natick, MA) using Psychophysics Toolbox (Brainard, 1997; Kleiner et al., 2007). Participants viewed stimuli from a distance of 248 cm on a monitor with a resolution of 1024 × 768 and a 100 Hz refresh rate. Responses were given on a keyboard.

Each trial consisted in the presentation of a randomly selected target morph for 500 ms, followed by a 1000 ms black-and-white pixel noise mask, intended to minimize afterimages. Participants then had to adjust a new morph to match the target expression. The initial adjustment morph was chosen randomly from the 147 morphs. Participants pressed the keyboard arrow keys to cycle through the morphs and pressed on the space bar when they had found the matching morph. We did not specify which feature participants should rely on to make their decisions and they were given unlimited time to respond (M=6 seconds, SD=2.4 seconds). After participants gave their response, a noise mask appeared on the screen for 1000 ms. The response morph was recorded as the numerical value of the selected morph, which ranged from 1 to 147. Participants each completed 588 trials in four blocks of 147 trials, each separated by a short break.

### EEG acquisition and preprocessing

EEG signals were acquired using a Brainvision cap with no online filtering. We used 128 Ag/AgCl electrodes mounted on a fabric cap and amplified by an ActiCHamp Plus amplifier (Brain Products) at a sampling rate of 1000 Hz. Data were referenced online to a unilateral electrode placed behind the left ear and the ground electrode was placed on the forehead. Electrodes were arranged according to the international 10–20 system.

Data preprocessing was performed using the FieldTrip toolbox (Oostenveld et al., 2010) implemented in MATLAB (The MathWorks, Natick, MA). Signals were re-referenced to the right mastoid electrode (M2) and subsequently high-pass filtered at 0.4 Hz using a finite impulse response (FIR) windowed-sinc filter (order=8000, two-pass). A detrending procedure was also applied to remove slow drifts from the continuous recordings. Data were then segmented into epochs time-locked to the onset of the target morph spanning 500 ms pre-stimulus to 1500 ms post-stimulus. Baseline correction was applied to segmented data using the interval –300 ms to 0 ms relative to stimulus onset. Finally, the data were low-pass filtered using a zero-phase Butterworth low-pass filter with a 100 Hz cutoff. Power-line noise was also attenuated using a discrete Fourier transform (DFT) filter at 50 Hz.

### Data analyses

We quantified serial dependence with response error, the shortest numerical distance between the target morph and the response morph. Error is thus given in morph units. This error was examined as a function of the previous morph such that its sign indicates the direction of error relative to the previous stimulus: positive error is attraction towards the previous stimulus whereas negative error is repulsion. Using a jackknife procedure, we removed trials in which response time and/or error were ±2SD from the mean calculated without that trial. We pooled trials across subjects.

Two types of consecutive trials exist. Consecutive morphs could either belong to the same emotion category (within-category trials) or not (between-category trials). Because of these two trial types, serial effects could be confused with prototype attraction. For within-category trials, there is a higher proportion of trials where the previous stimulus was towards the prototype than away from it. Thus if participants respond that the current stimulus is more like the previous stimulus than expected by chance, this could be due to an attraction to the previous or to the prototype. To overcome this confusion, we fit a sinusoid function to the relationship between error and morph units, and performed all other analyses on the residuals (see Pascucci et al., 2019; Pascucci et al., 2023 for a similar approach).

### Representational Similarity Analysis

RSA is a correlational approach that quantifies the geometry of our three measured variables: perceptual similarity, serial dependence and EEG. The perceptual similarity matrix was obtained in the preliminary experiment. The serial dependence matrix was a 147×147 matrix in which each cell *(i*,*j)* contained the response error when morph *i* was preceded by morph *j*. We averaged cells *(i*,*j)* with cells *(j*,*i)* such that the matrix was symmetric around the diagonal. We interpolated missing pairs by replacing missing values with the average of the 8 closest neighbors, with wrap-around indexing at matrix boundaries. We smoothed the resulting matrix by averaging across a 5-cell-radius neighborhood. This smoothing was used for all data matrices described hereafter.

We generated an EEG_n_ matrix by correlating the brain activity evoked by the presentation of a morph *i* with the activity evoked by the presentation of a morph *j*. Because EEG is time-resolved, there is actually a matrix for each recorded time point. In order to investigate the temporal pattern of brain activity, we created 18 matrices corresponding to 18 100-ms long time bins (3 pre-stimulus and 15 post-stimulus). For each morph, time bin, and electrode, we computed average amplitude. We then correlated that with the average obtained for another morph.

We generated an EEG_n-1_ matrix to assess how prior stimulus history influences subsequent neural responses. The brain activity evoked by the presentation of a morph *i* was correlated with the activity from all trials that were preceded by morph *j*. Again, we generated 18 EEG matrices corresponding to 18 100-ms time bins across all electrodes. The EEG_n-1_ matrices were non-symmetrical.

We correlated perceptual similarity, serial dependence and EEG matrices with five theoretical models (Figure 3). The first model expresses the physical difference between pairs of morphs in morph units (Fig 3a). This model was used to describe low-level features; given that the morphs were created by a linear variation of the percentage of prototypes, we expected our model to vary low-level features linearly. Nevertheless, as a better measure of how low-level features vary throughout the morph continuum, we quantified the composition of the morph images with a Gabor filterbank algorithm (Fig 3b; Leeds et al., 2013). Each morph image is passed through a bank of Gabor filters that vary in orientation, spatial scale, and position. The filters approximate the way primary visual cortex neurons encode visual information, yielding a V1-like representation of the image. Similarity between images is then computed using the euclidean distance between vectors of each stimulus. The third model captures categorical perception (Fig 3c). To create this model, we started with the physical similarity matrix (in morph units) and divided the similarity value by 1.5 when two morphs did not belong to the same category. Given that we use a correlational approach, the value of 1.5 is arbitrary, and only the resulting shape of the model matrix counts. The fourth model expresses the emotional ambiguity of the stimulus (Fig 3d). We reasoned that near-prototype morphs are more emotionally salient, whereas morphs near category boundaries are ambiguous and less salient. We started with the morph unit similarity matrix and divided the similarity value by 4 for pairs in which at least one was ambiguous (defined as within a 5 morph-unit radius from a category boundary). The fifth model aimed to capture what we called a full-feature-tuning pattern (Fig 3e). Serial dependence is feature tuned (Manassi et al., 2023): it occurs most strongly between stimuli that are not too different from each other. When two consecutive stimuli are very dissimilar, serial dependence is small or null. Serial dependence is also reduced when two consecutive stimuli are too similar, simply because the stimulus space in which such effects could operate is itself reduced. This explains the use of a derivative of Gaussian to characterize the pattern of serial dependence. The term “full” feature pattern refers to the decrease of SD when consecutive stimuli are *both* too similar and too dissimilar. We created a matrix to model this pattern, that also included the categorical model. We started with the categorical model (model #3) to which we added feature-tuning by dividing the value of a cell by 5 if the pair of morphs was very similar (defined as over 80% of the maximum value of the categorical matrix) or very different (<15%). These five theoretical models allowed us to investigate the geometry of our three measured variables: perceptual similarity, behavioral serial dependence, and neural measures. We also correlated the measures amongst themselves.

**Figure 3.**
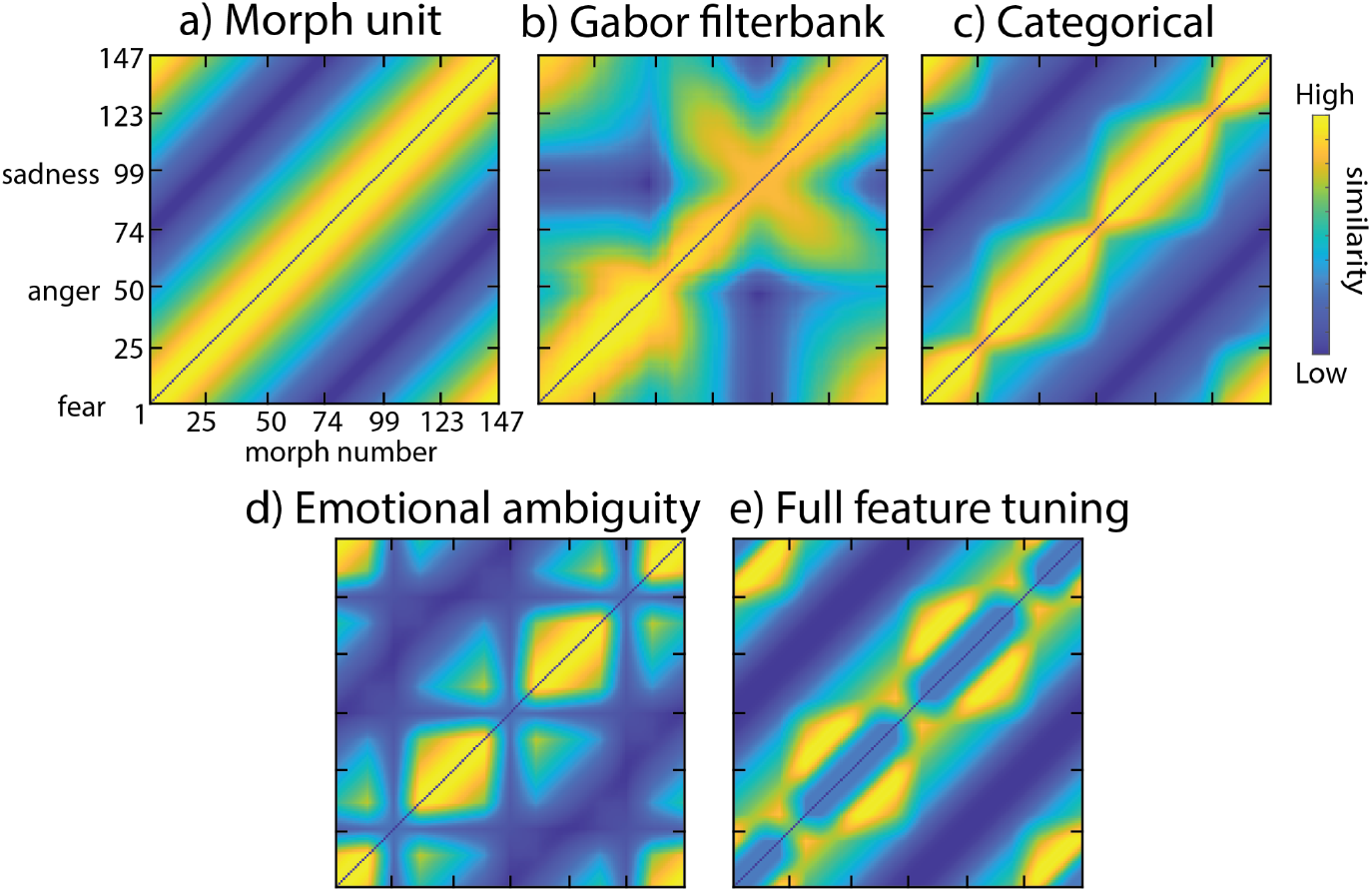
Representational similarity matrices for each model across the stimulus set. Similarity units are in (a) morph units; (b) Gabor filterbank units; (c)-(e) arbitrary units.

We performed Pearson correlations between matrices; given that we were not interested in the linear combination of several matrices to explain the geometry of another, collinearity was not an issue and we simply compared Pearson correlation coefficients. We ascertained significance by performing permutation tests. For example, to question the difference (delta) between two correlation coefficients resulting from the correlation of matrices A and B versus A and C, we shuffled the values of B (and C) cells and calculated two new correlations with A, and a delta between them. By permuting 1000 times, we calculated a null distribution of the delta between the two correlation coefficients.

Our hypotheses were the following: (1) We expected the perceptual similarity matrix to correlate the most with the categorical model. (2) For the serial dependence matrix, our hypothesis was of a higher correlation with the full-feature-tuning model matrix than with the other models. Indeed, we hypothesized that SD occurs at a high representational level, implying a categorical effect as well as feature tuning. (4) Concerning the neural representation, we expected low-level features to dominate the neural signal at early stages of processing. At later stages, the neural signal may reflect the perceptual representation or subjective experience of the stimulus. Accordingly, EEG matrices derived from later timepoints may correlate with matrices indexing higher-level processing, such as the perceptual similarity matrix or the categorical model (its less noisy theoretical counterpart). (5) Finally, looking at the EEG activity that may express a trace of the past in the response to the current stimulus, we hypothesized that this trace would represent high-level characteristics. We did not have any strong prior expectations regarding the specific time point at which such a pattern might emerge.

### Classic Serial Dependence Analysis

To facilitate comparison with the literature, we also quantified serial dependence by examining response error as a function of the relative difference between current and previous target morphs. This relative difference between consecutive morphs was defined in three ways. First, in terms of morph units (the shortest numerical distance between the current and previous morphs). Second, in terms of low-level feature similarity as estimated by the Gabor filterbank algorithm. Third, in terms of the perceived difference between consecutive morphs, as determined by the preliminary experiment, i.e. categorically. In all cases, we smoothed the data by averaging across bins of relative differences: from minimum to maximum, with 14 bins evenly spaced. To investigate differences between the within- and between-category trials, we fitted the data with specific functions and used permutation tests to assess significance. For relative differences between consecutive morphs expressed as morph units and Gabor filterbank algorithm estimates, we fitted with exponentials weighted by the standard error of the mean. We then ran a permutation test to evaluate whether the intercept and slope differed between conditions. To do so, we randomized the mapping between relative difference and response errors, fitted an exponential function to the shuffled data, and took the resulting differences between within- and between-category conditions across a thousand iterations to build a null distribution. For the relative perceived difference, we fitted both conditions with a weighted Gaussian and ran the same permutation test on the amplitude of the fitted curves.

Consistent with our hypothesis, we expected SD feature tuning to differ depending on whether differences between stimuli were expressed in physical versus categorical terms. When expressed on a physical scale, we expected an exponential pattern of response errors and greater serial dependence for equivalent physical distances when a pair was between-category than within-category. When differences between morphs were expressed in perceptual terms, we expected a Gaussian pattern of response errors, with no difference between the within- and between-category pairs. Indeed, for a given low-level difference between morphs, the perceived difference depends on their categorical membership. Hence, if serial dependence operates at a categorical level, we should observe a difference in response errors between the two conditions for the same low-level difference.

## RESULTS

### Preliminary experiment

The preliminary experiment showed that our stimuli were perceived categorically. Two analyses support this hypothesis.

First, we correlated the perceptual similarity matrix with four models to characterize its geometry (Fig 4a-b). Recall that each cell in the similarity matrix corresponds to the number of times the pair was unselected together per number of times the pair was presented. The categorical aspect of the matrix can already be seen in the blobs of higher similarity around the three prototypes, and lower similarity between them, corresponding to category boundaries. The perceived similarity matrix correlated most with the categorical model (rho=0.90, permutation test p-value<0.0001). It also correlated, but to a lesser extent, with the morph unit model (rho=0.82, p-value<0.0001; difference with categorical=0.08, permutation test p<0.0001), the Gabor filterbank model (rho=0.72, p<0.0001; difference with categorical=0.18, p<0.0001) and the emotional ambiguity model (rho=0.71, p<0.0001; difference with categorical=0.19, p<0.0001).

**Figure 4.**
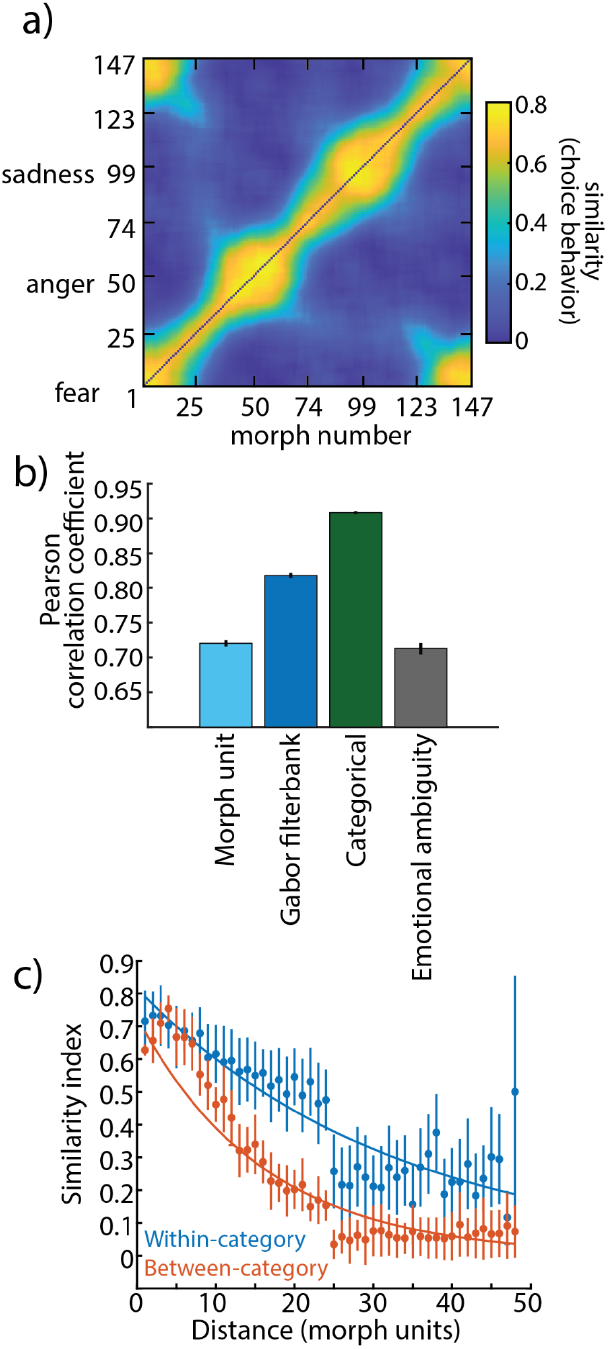
a) Perceptual similarity matrix. Each cell contains the similarity between a morph pair. b) Pearson correlation coefficient between the perceived similarity matrix and model matrices (morph unit, Gabor filterbank, categorical and emotional ambiguity). Error bars are bootstrapped 95% confidence intervals. c) Mean similarity index as a function of morph-units distance between morph pairs. Error bars indicate standard deviation. Curves represent weighted exponential functions fit separately for within-versus between-category trials.

Second, we examined perceived similarity as a function of the difference, in morph units, between pairs of stimuli, separately for pairs belonging to the same category (within-category) or not (between-categories) (Fig 4c). Similarity decreased with increasing distance, but for a given distance, perceived similarity was greater when two stimuli belonged to the same category than when they did not. This is supported by the fact that the intercept of the two fitted exponential functions were significantly different (within-category intercept=0.82; between-category intercept=0.73; delta=0.09; p<0.0001).

## Main experiment

### Task performance

Participants reported the perceived emotion by adjusting a response morph to match the target morph (Fig. 2). Figure 5 presents behavioral performance. Presented morph numbers are around the outer circumference of the circle, at the origins of the colored lines, and are therefore evenly spaced. Average matching responses are points on the inner circumference, such that any lines that are not perpendicular to the outer circumference indicate deviations from veridical responses. The clusters of points around the fear, anger and sadness prototypes show the classic prototype attraction effect. The data in Figure 5 is uncorrected for this prototype effect. All subsequent analyses work on the residuals of a function fit to the relationship between error and morph units, which unconfounds the prototype effect from serial dependence (see Methods).

**Figure 5.**
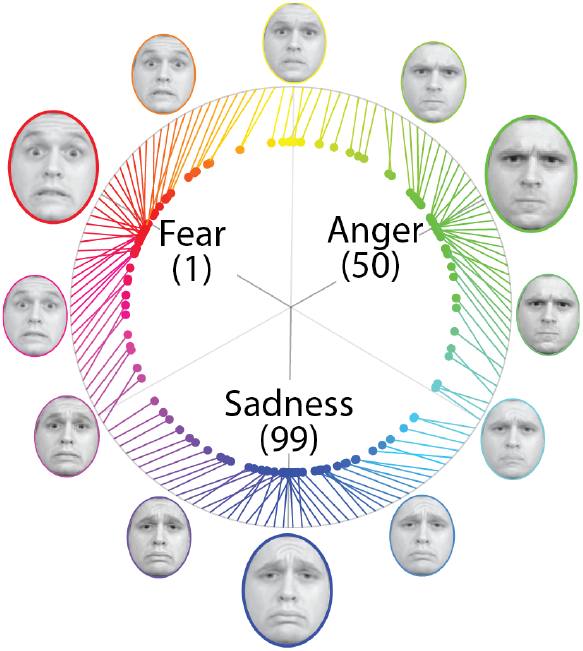
Behavioral performance in the main experiment. The colored lines originate at morph number, and their endpoint is located at the perceived morph number. The three prototypes: fear (morph #1), anger (morph #50) and sadness (morph #99), were taken from the Radboud Face Database (RaFD).

### Representational Similarity Analysis

We correlated the amplitude evoked by the presentation of one stimulus across all electrodes with the amplitude evoked by each other stimulus. The resulting matrices, for each 100-ms time-window, were then correlated with the empirical and theoretical models of interest (Fig 6). The correlation with the emotional ambiguity model was the highest, showing a pronounced and sustained elevation between 200 and 700 ms, during which it reached the highest correlation of all models (in the 400-500ms window). Conversely, the correlation with the Gabor filterbank model was the weakest, with a strong dip around 200 ms and generally low or negative values across the rest of the time course. The correlations with the three remaining models (morph unit, perceptual similarity, categorical) were highly similar across the entire epoch. They showed a peak at stimulus onset, a dip shortly after (in the 100–200 ms window), a secondary rise peaking around 500–600 ms, and a gradual decline toward baseline in the later portion of the time course.

**Figure 6.**
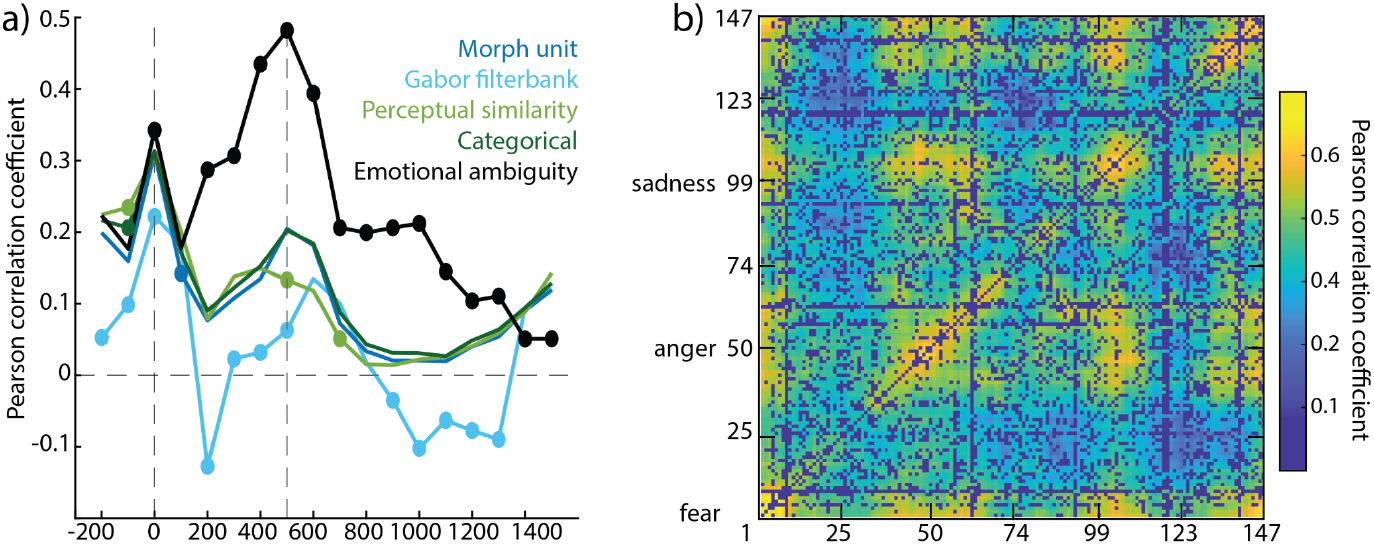
a) Pearson correlation coefficients for the correlation of the EEG matrix and model matrices (morph unit, Gabor filterbank, perceptual similarity, categorical, and emotional ambiguity), for each 100-ms time window. Vertical dashed lines indicate stimulus onset and offset. Filled circles denote time points at which the correlation coefficient for one model differed significantly from all others (permutation test, α=0.05). b) EEG matrix for the 400-500ms time window (in which there was the maximal correlation with the emotional ambiguity model). Each cell *(i*,*j)* contains the coefficient of the correlation between the activity evoked by morph *i* and the activity evoked by morph *j* (averaged with cell *(j*,*i)*).

Serial dependence is represented as a matrix in Fig 7a. Each cell expressed the error in the matching task for a given morph *i* preceded by another morph *j* (averaged over the diagonal). This SD matrix correlated the most with the full-feature-tuning model (rho=0.29, p<0.0001) and, to a lesser extent, with the morph unit (rho=0.23, p<0.0001), categorical (rho=0.14, p<0.0001), perceptual similarity (rho=0.07, p<0.0001), Gabor filterbank (rho=0.05, p<0.0001) and emotional ambiguity models (rho=0.04, p<0.0001) (Fig 7b). All coefficients were different from each other (ps<0.0001), except the correlation with Gabor filterbank vs with perceptual similarity (delta=0.02, p=0.07) and with Gabor filterbank vs with emotional ambiguity (delta=0.01, p=0.11).

**Figure 7.**
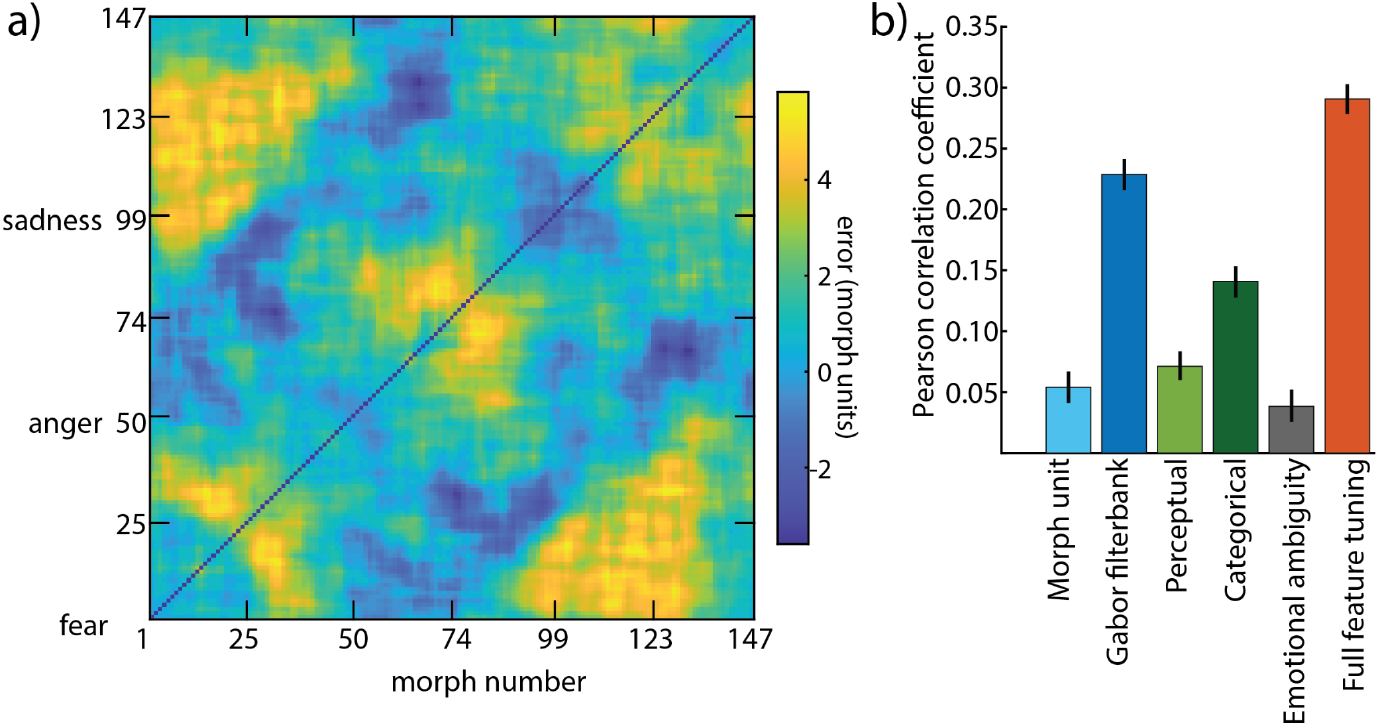
a) Serial dependence matrix. Each cell *(i*,*j)* corresponds to the error, in morph units, between current morph *i* and previous morph *j*. b) Pearson correlation coefficient between the serial dependence matrix and model matrices (morph unit, Gabor filterbank, perceptual, categorical, emotional ambiguity and full feature-tuning). Error bars indicate 95% confidence intervals estimated via permutation testing.

Finally, we described the neural representation of the previous stimulus by correlating the activity evoked by a morph *i* with the activity evoked by trials that were preceded by each morph *j*, again in 100-ms time windows. Rows in the matrix correspond to current morphs, and columns to previous morphs. All matrices were then correlated with the empirical and model matrices (Fig 8). The correlation with the emotional ambiguity model was the strongest up to approximately 1300 ms following stimulus onset, reaching a maximum in the 400–700 ms time window. In contrast, there was a pronounced negative correlation with behavioral SD, and with the full feature-tuning model. The remaining models (Gabor filterbank, morph unit, categorical, and perceptual similarity) correlated minimally with the EEG_n-1_ matrix, with correlation values that were weakly positive or not significantly different from zero.

**Figure 8.**
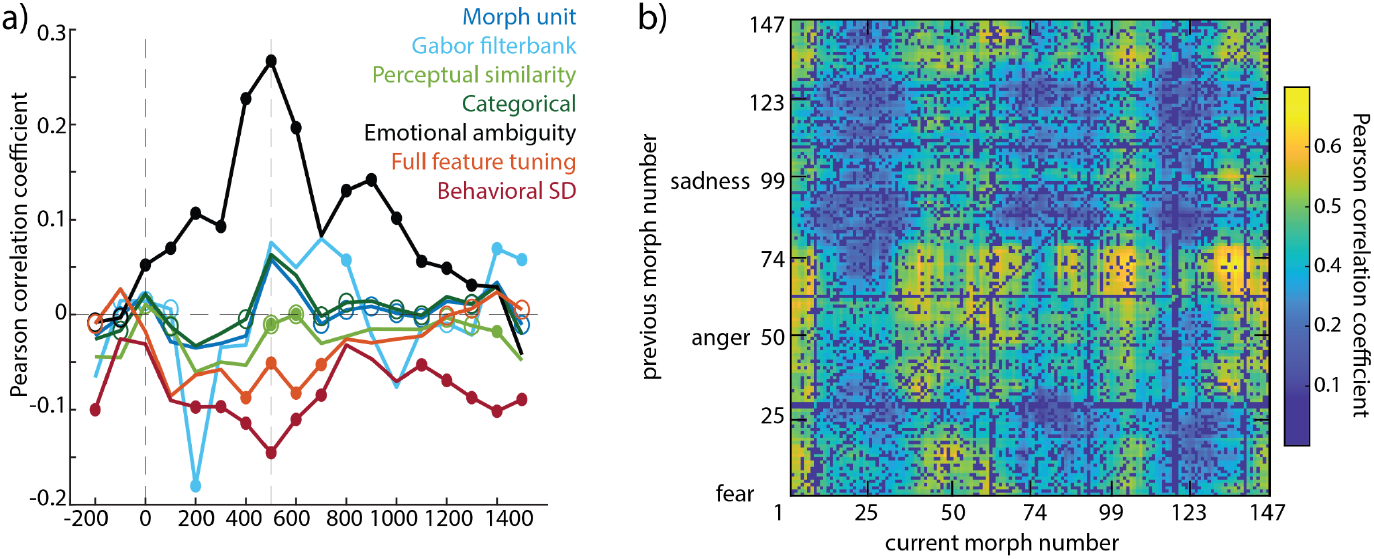
a) Pearson correlation coefficients between the EEG_n-1_ matrix and each model matrix (morph unit, Gabor filterbank, perceptual similarity, categorical, emotional ambiguity, full feature-tuning, behavioral serial dependence), at each 100-ms time point. Conventions as in Figure 6a, here, empty circles denote time points at which the correlation coefficient for one model does not differ significantly from zero (permutation test, α = 0.05). b) EEG_n-1_ matrix for the 400-500ms time window (in which there is the maximal correlation with the emotional ambiguity model). Each cell *(i*,*j)* contains the coefficient of the correlation between the activity evoked by morph *i* and the activity evoked by all morphs preceded by *j*.

### Classic Analysis

In a second approach, closer to what is done in the literature, we quantified serial dependence by examining response error as a function of the difference between two consecutive stimuli. We defined the relative difference between consecutive stimuli in three ways.

The first expressed relative difference in morph units (Fig 9a). We observed a negative exponential pattern for the between-category trials (slope=-0.03; p<0.0001) and no significant slope for the within-category trials (0.009; p=0.09). This absence of a significant slope suggests that all relative within-category differences were equally able to induce serial dependence, although descriptively there was a drop off for larger relative differences. A difference between the two conditions appears for short distances, with between-category trials eliciting a stronger attractive bias than within-category trials. This is supported by a significant difference in the intercept of the exponential fits (delta=2.55; p < 0.0001).

**Figure 9.**
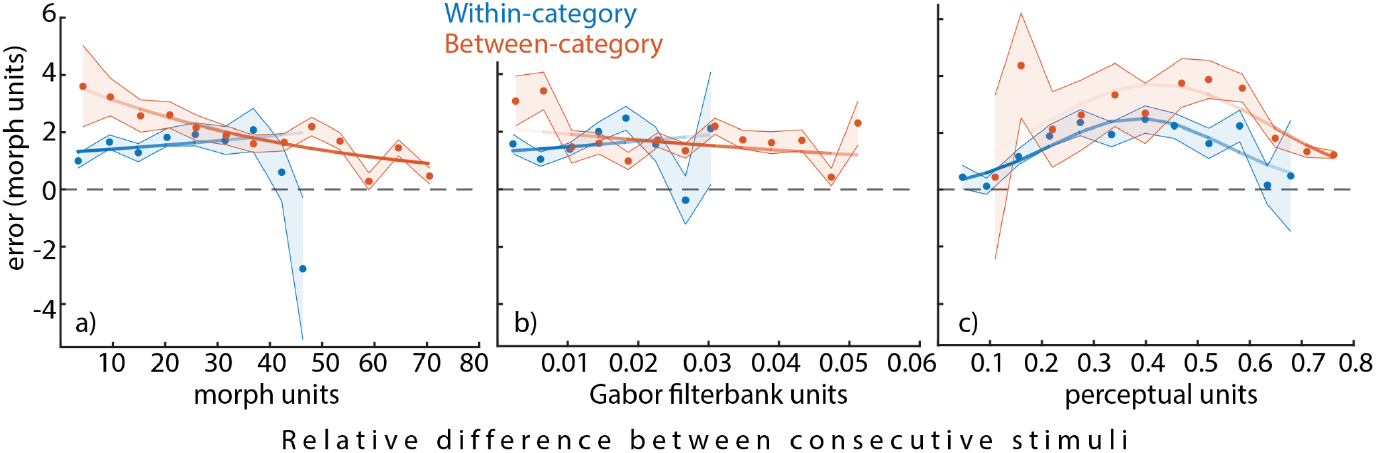
Mean error as a function of the relative difference between consecutive stimuli for within- and between-category trials. Shaded areas represent standard error of the mean, transparency represents the amount of data that contributed to each point. a) Relative difference in morph units. b) Relative difference in gabor filterbank arbitrary units. c) Relative difference in perceptual similarity, based on choice behavior.

The second way we quantified the relative difference between consecutive stimuli was to use dissimilarity as computed by the Gabor filterbank algorithm. Again, for each bin of relative difference, we calculated mean response error (Fig 9b). We found a negative exponential pattern for the between-category trials (slope=-11.5; p=0.03) and no significant slope for the within-category trials (slope=12.4; p=0.16). Again, the difference appeared for short relative distances, with greater error for between-category trials, supported by significantly different intercepts of the exponential fits (delta=0.85; p=0.02).

These first two analyses express differences between consecutive stimuli in physical units, and support the idea that when two consecutive stimuli do not belong to the same category, the serial attractive effect is stronger for short distances.

The third way we defined the relative difference between consecutive stimuli was as a perceived difference (based on similarity measured in the preliminary experiment; Fig 9c). Both within- and between-category trials followed a Gaussian pattern: error decreased when two consecutive stimuli were perceptually very similar or very dissimilar. For intermediate relative differences, there was an attractive effect: observers tended to err in the direction of previous stimuli. We fitted Gaussians; no amplitude difference was observed between the two conditions (delta=1.22; p=0.28).

## DISCUSSION

### The perception of emotional faces is neurally and psychologically categorical

The emotional expression of faces was perceived categorically. This means that small image variations within a category (for example, in the vicinity of a prototypically fearful face), were perceived to differ very little, whereas similar image variations across category boundaries (e.g. across the fearful versus sadness boundary) were perceived as highly different. Categorical perception of facial expressions, first reported by Etcoff et Magee (1992), has been replicated many times, including with representational similarity analysis similar to our approach (Brooks & Freeman, 2018; Murray et al., 2021). Other classic reports of categorical perception include speech sounds (Liberman et al., 1957) and color (Bornstein and Korda, 1984). Representational similarity analysis overcomes differences in format between different measures or scales (Kriegeskorte, 2008), and allowed us to bridge the gap between theoretical, behavioral and neural representations.

The *neural* representation of emotional faces also expressed high-level information, but in contrast to the categorical information contained in the mental representation, emotional ambiguity was predominant in the EEG response to faces. Indeed, there was a high correlation between near-prototypical face morphs, regardless of which prototype they were near. For example, a prototypically fearful face correlated highly with a prototypically sad face, and less with faces near category boundaries.

Many authors propose that the perception of emotions, in addition to being categorical, is also determined by two distinct dimensions: arousal (i.e. level of energy) and valence (i.e. hedonic tone) (Russel, 1980; Fujimura et al., 2011). These two determinants of perception are not independent (Baudouin et al., 2025). In the present study, the three emotions (anger, fear and sadness) share negative valence. We did not gather ratings of arousal for our face morphs, nor did we attempt to control the level of arousal between morphs. However, it may be the case that ambiguous morphs were less arousing than near-prototypical faces. Thus, the neural representation picked up by the EEG could be driven by brain areas that respond to arousal, such as the right fusiform face area (FFA) and left insula (Matsuda et al., 2013). Right FFA and other brain areas have been shown to respond differentially to prototypical emotional faces versus ambiguous ones (Matsuda et al., 2013; Wang et al., 2017). Unfortunately, we do not have enough data to perform a searchlight analysis that might shed light on different dimensions being preferentially expressed in different sub-groups of electrodes. Another difference between salient emotional faces and ambiguous between-category morphs is that salient emotions are responded to with higher confidence (Wang et al., 2017). Numerous studies have shown distinct neural signatures of high-versus low-confidence percepts (Gherman & Philiastides, 2014; Boldt & Yeung, 2015), and our results could very well be explained by this confidence difference.

Although the emotional ambiguity model correlated best with the neural representation, categorical, perceptual similarity and morph-unit models also correlated with the EEG response. The fact that multiple models correlated with brain activity is not surprising, given that these models were also correlated amongst themselves. The relative strength of each model may relate to the fact that different brain areas express different dimensions of the faces, and these activities become predominant at different times during face processing.

### Serial dependence of facial expression was categorically feature tuned

We observed attractive serial dependence, replicating multiple other studies on facial expression (Alais et al., 2021; Collins, 2022; A. Liberman et al., 2014, 2018). Feature tuning is a well documented property of serial dependence (Manassi et al., 2023): a past stimulus influences current perception only if the feature distance between previous and current stimuli is neither too small nor not too large. Indeed, small inter-stimulus distances inherently constrain the measurable magnitude of an attractive effect, i.e. there is little space in which an identical past could influence the perception of the current stimulus. For large distances, the assumption that the two samples represent the same object is likely broken, interrupting temporal integration (Collins, 2022). The current study shows that feature tuning operates on a categorical scale: the strength of serial dependence between two stimuli is determined by their distance in a perceptual space shaped by category membership.

The model that expresses these two key characteristics was the full feature-tuning model. Indeed, this model combines feature tuning in physical image units with categorical perception, and it correlated most with the pattern of serial dependence observed across morph pairs. We also adopted a more classic analytical approach to show that serial dependence is tuned to perceived differences rather than physical differences. We examined serial dependence as a function of the distance between consecutive stimuli. When that distance was defined in physical units (morph or Gabor filterbank units), there was more serial dependence between consecutive between-category morphs than between consecutive within-category morphs. When we defined distance in perceptual units, taking into account categories, there was no longer a difference between within-versus between-category pairs and only feature tuning remained. This pattern of results between physical distance versus perceptual-categorical distance makes sense because small physical distances across category boundaries are perceptually large and appropriate to cause serial dependence. Small physical distances within a category correspond to very small perceived distances, too small to elicit much serial dependence. Therefore, realigning physical distances into perceptual-categorical space equalizes within- and between-category conditions and recovers canonical feature tuning. This idea is illustrated in Figure 10.

**Figure 10.**
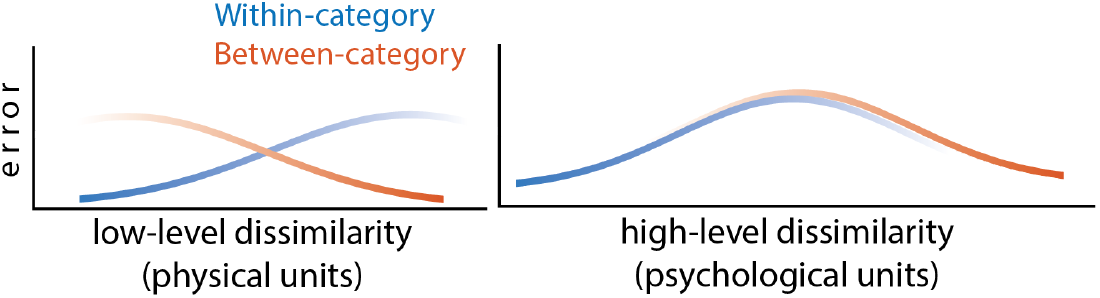
A visual explanation of our interpretation. The left-hand panel schematizes the observed error as a function of low-level dissimilarity, expressed in physical units (like morph units or gabor filterbank units). Small low-level dissimilarity leads to high error for between-category morph pairs and low error for within-category morph pairs. The color gradient represents the amount of data, with darker colors meaning more data (e.g. there should be few between-category pairs that are close in image space). When the dissimilarity between these same pairs is expressed in psychological units (right panel), the two curves realign.

One limitation of our study concerns the lack of strict control of the low-level features of our face morphs. Morph units are not a real physical property and result from experimental choices. To tackle this issue, we included an estimate of the physical features of our face morphs by using the gabor filterbank algorithm to quantify low-level features. The resulting representational matrix differed from the morph-unit model. However, neither behavioral measures of serial dependence nor neural representation analyses revealed differences between these two physical scales. Another limitation comes from the fact that the estimate of the perceptual representation of our morphs was obtained from a group of participants different from those who performed the main experiment. Future work could benefit from assessing perceptual representations for each participant, allowing the estimation of individual category boundaries and the investigation of inter-individual differences in the feature tuning of serial dependence.

### Implications for the source and site of serial dependence

The source of serial dependence refers to the representation of the past that influences current perception. The site of serial dependence refers to the information about the current stimulus that is influenced by this past representation. Both the source and site could be at different levels of processing: either low-level representations of physical stimulus features, or high-level representations that include context, categories, or other interpretations. Several authors have shown that source and site may be context- or stimulus-dependent.

For example, Cicchini et al. (2021) used the surround tilt illusion, in which the perceived orientation of a Gabor is repulsed from a tilted visual context. They demonstrated that when a Gabor presented in a neutral context was preceded by a Gabor presented in a surround-tilt context, the perceived orientation of the current Gabor was attracted to the previous illusory orientation rather than to the previous physical orientation. They concluded that the source of serial dependence, i.e. the prior that influenced current percept, was generated at a high representational level because it included the context effect. When a Gabor presented in a surround-tilt context was preceded by a neutral Gabor, the current Gabor was (of course) perceived with an illusory tilt, but only after it had been influenced by the physical orientation of the previous neutral Gabor. Thus, while the source of serial dependence was a high-level representation, the site of serial dependence was at the level of a low-level feature representation.

Similarly, Collins (2022) showed that serial dependence can occur on high- or low-level representations, this time depending on the type of stimulus. Arbitrary shapes made up of conjunctions of visual features were presented, and participants reported shape. Shape percepts were serially dependent, and slightly more so when other object features matched between consecutive stimuli. Shape perception is likely low level and the representation of past shape that influences current shape perception is only minimally determined by its integration into an object file. However, serial dependence of emotional expression occurred robustly between consecutive faces of the same identity, but was reduced when the consecutive faces had a different identity. The perception of emotional face expression is likely high level and the representation of past expressions that influences current emotion perception is highly determined by its interpretation (as a distinct individual).

Moreover, a study by Sheehan et al. (2024) suggests that serial dependence can be mediated by a prior shared across different neural circuits. Indeed, they showed an attractive bias towards a previously imagined stimulus, in the absence of a physically presented stimulus. Similar to this stimulus-dependency (also in Collins, 2022) and context-dependency (Cicinni et al., 2021), the current study offers new evidence in favor of a high-level representation as the source of serial dependence. We believe that this supports the notion that when the interpretation of a stimulus requires high level representations, as is likely the case for facial emotions, the feature tuning of serial dependence aligns with that representational level. In other words, the temporal integration of visual information appears to be guided by the perceptually relevant mental representation of the stimulus, rather than by its physical properties alone. Our design does not allow us to draw conclusions about the site at which serial dependence operates. Future work could examine to what extent particular task representations determine the temporal integration of sensory information. In the context of face perception, high-level properties such as facial emotion may influence the perception of low-level features (e.g. orientation of the eyebrows) and serial dependence of those features.

In conclusion, we argue that temporal integration of sensory information depends on the relevant representation for a particular task.

### Neural representation of the past

We observed the trace of a past stimulus in the response to the current stimulus. This trace was categorical: the response evoked by an emotionally unambiguous face was similar to the trace of an emotionally unambiguous face, and dissimilar to ambiguous faces. This means that the neural trace of the previous stimulus carried information about emotional ambiguity. We show here that the processing of the current stimulus carries information about the past, replicating other studies on serial dependence (Bae & Luck, 2019; Collins et al., 2024; Luo & Collins, 2023; Ranieri et al., 2022). Interestingly, the lasting previous information – or reactivated information – appeared to convey the same high-level content as the neural signals associated with current processing.

The neural pattern was anti-correlated with the geometry of behavioral serial dependence. Sheehan & Serences (2022) found a similar pattern of anti-correlated behavior and fMRI activity evoked by oriented Gabor patches. There was attractive serial dependence between successive orientations, but adaptation of the fMRI response such that activity appeared repulsed from the previous orientation. They modeled their results by assuming that serial dependence arises in post-sensory brain areas that take adaptation into account when decoding incoming (adapted) sensory signals. Of course, the meaning of our anti-correlation is very different from theirs. In the context of EEG representational similarity analysis, our anti-correlation means that the stimulus values that lead to the most behavioral error in the direction of the previous stimulus are the values for which the representation of the past is least like the representation of current stimuli.

The behavioral serial dependence matrix shows the morphs that are most likely to succumb to and cause serial dependence. Some morphs are therefore more susceptible than others, and it is precisely those morphs whose mnemonic representation differs most from the sensory representation. Our results therefore argue for a dissociation between sensory and memory representations, and in this respect our results align closely with several other investigations of the brain correlates of serial dependence (Sheehan & Serences, 2022; Luo & Collins, 2023; Hajonides et al., 2023).

